# Integrated genomic view of SARS-CoV-2 in India

**DOI:** 10.1101/2020.06.04.128751

**Authors:** Pramod Kumar, Rajesh Pandey, Pooja Sharma, Mahesh S Dhar, A Vivekanand, Bharathram Uppili, Himanshu Vashisht, Saruchi Wadhwa, Nishu Tyagi, Saman Fatihi, Uma Sharma, Priyanka Singh, Hemlata Lall, Meena Datta, Poonam Gupta, Nidhi Saini, Aarti Tewari, Bibhash Nandi, Dhirendra Kumar, Satyabrata Bag, Deepanshi, Surabhi Rathore, Nidhi Jatana, Varun Jaiswal, Hema Gogia, Preeti Madan, Simrita Singh, Prateek Singh, Debasis Dash, Manju Bala, Sandhya Kabra, Sujeet Singh, Mitali Mukerji, Lipi Thukral, Mohammed Faruq, Anurag Agrawal, Partha Rakshit

**Affiliations:** National Centre for Disease Control (NCDC), 22-Shamnath Marg, Delhi-110054, India; CSIR-Institute of Genomics and Integrative Biology (CSIR-IGIB), Mall Road, Delhi-110007, India

**Keywords:** COVID-19, SARS-CoV-2, Whole Genome Sequencing (WGS), MinION and Phylogenetics

## Abstract

India first detected SARS-CoV-2, causal agent of COVID-19 in late January-2020, imported from Wuhan, China. March-2020 onwards; importation of cases from rest of the countries followed by seeding of local transmission triggered further outbreaks in India. We used ARTIC protocol based tiling amplicon sequencing of SARS-CoV-2 (n=104) from different states of India using a combination of MinION and MinIT from Oxford Nanopore Technology to understand introduction and local transmission. The analyses revealed multiple introductions of SARS-CoV-2 from Europe and Asia following local transmission. The most prevalent genomes with patterns of variance (confined in a cluster) remain unclassified, here, proposed as A4-clade based on its divergence within A-cluster. The viral haplotypes may link their persistence to geo-climatic conditions and host response. Despite the effectiveness of non-therapeutic interventions in India, multipronged strategies including molecular surveillance based on real-time viral genomic data is of paramount importance for a timely management of the pandemic.

## INTRODUCTION

The rapid worldwide spread of a novel coronavirus following its first appearance in China has pressed the global community to take measures to flatten its transmission (Chan et al., 2020; Zhu et al., 2020). The virus was named as “SARS-CoV-2” and the disease it causes has been defined as “coronavirus disease 2019” (COVID-19) (CSG, 2020). WHO declared COVID-19 a Pandemic on 11^th^ of March, 2020 based on the speed and scale of transmission with almost 4.7 million cases reported across the globe, so far (WHO, 18th May 2020). The common signs of infection by SARS-CoV-2 include cough, fever, sore throat, respiratory symptoms inclusive of shortness/difficulties in breathing. More severe symptoms can include pneumonia, severe acute respiratory syndrome, kidney failure and even death with coalescence of factors (Zhu et al., 2020, Youg et al., 2020). Many COVID-19 cases have been reported to be asymptomatic and may serve as carrier of SARS-CoV-2 (Xu et al., 2020; He et al., 2020). Whole Genome Sequences (WGS) of SARS-CoV-2 suggest bat-CoV to be its closest progenitor with 96% homology but, the RNA binding domain (RBD) of its spike protein with an efficient binding to ACE-2, the receptor for SARS-CoV-2 in human cell seems to have been derived from Pangolin-CoVs (Wan et al., 2020, Andersen et al., 2020). Next generation sequencing (NGS) aided understanding of evolution of SARS-CoV-2 genomes and its transmission patterns after it enters a new population is proving to be an important step towards formulating strategies for management of this pandemic (Chen and Li et al., 2020).

The first three cases in India were reported from the state of Kerala in late January and early February, with a travel history of Wuhan, China. India took drastic steps to contain the further spread of the virus including imposition of travel restrictions to-and-from the affected countries. There were no new cases of COVID-19 for almost a month. All three cases subsequently tested negative making India free of the disease at that point of time (Press Information Bureau, PIB, India, 2020). However, while the global focus was on China and other eastern countries like South Korea and Japan; European countries, middle-east and the USA reported a surge in cases of COVID-19, pressing the WHO to declare it as a pandemic. March 2020 onwards, India also witnessed a surge of imported cases from countries other than China which has been further assisted with local transmission. In March, imposition of nationwide lockdown checked the epidemic curve. Despite these measurements, India is at the verge of a large outbreak as the transmission is rapidly increasing with more than 100,000 cases of COVID-19 having been reported in the third week of May, 2020.

We carried out WGS of SARS-CoV-2 (n=104) from Pan-India through the network of Integrated Disease Surveillance Program (IDSP) of National Centre of Disease Control (NCDC), Delhi. We report here a comprehensive and integrative genomic view of SARS-CoV-2 in the Indian subcontinent. In this study, we combine genetic and epidemiological data to understand the genetic diversity, evolution, and epidemiology of SARS-CoV-2 across India. The spectrum of variations would be an important tool towards contact tracing, effective diagnostics and backbone for drug and vaccine development.

## METHODS

### Subject recruitment

The study was conducted jointly by the NCDC and CSIR-Institute of Genomics and Integrative Biology (CSIR-IGIB). Institutional ethical clearance was obtained at both the places prior to initiation of research. A total of 127 laboratory confirmed cases of COVID-19 from a targeted testing representing different locations (as described in **Table-1** and **supplementary figure-1**) were included in the study for genomic analyses. Targeted testing involved suspected cases; having symptoms (fever, cough and breathlessness) with recent travel history to high-risk countries or positive contacts of COVID-19 cases.

**Table-1:**
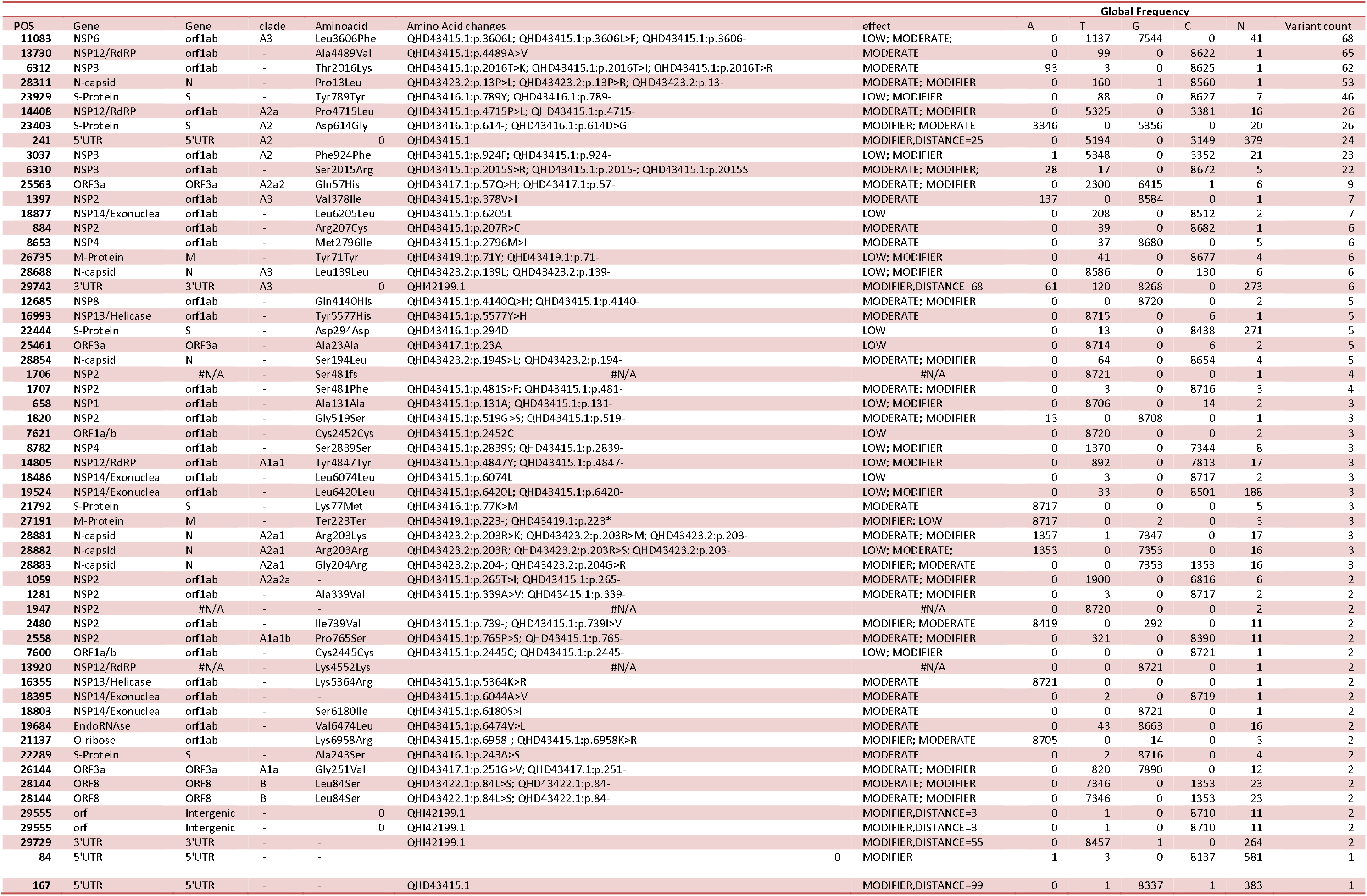
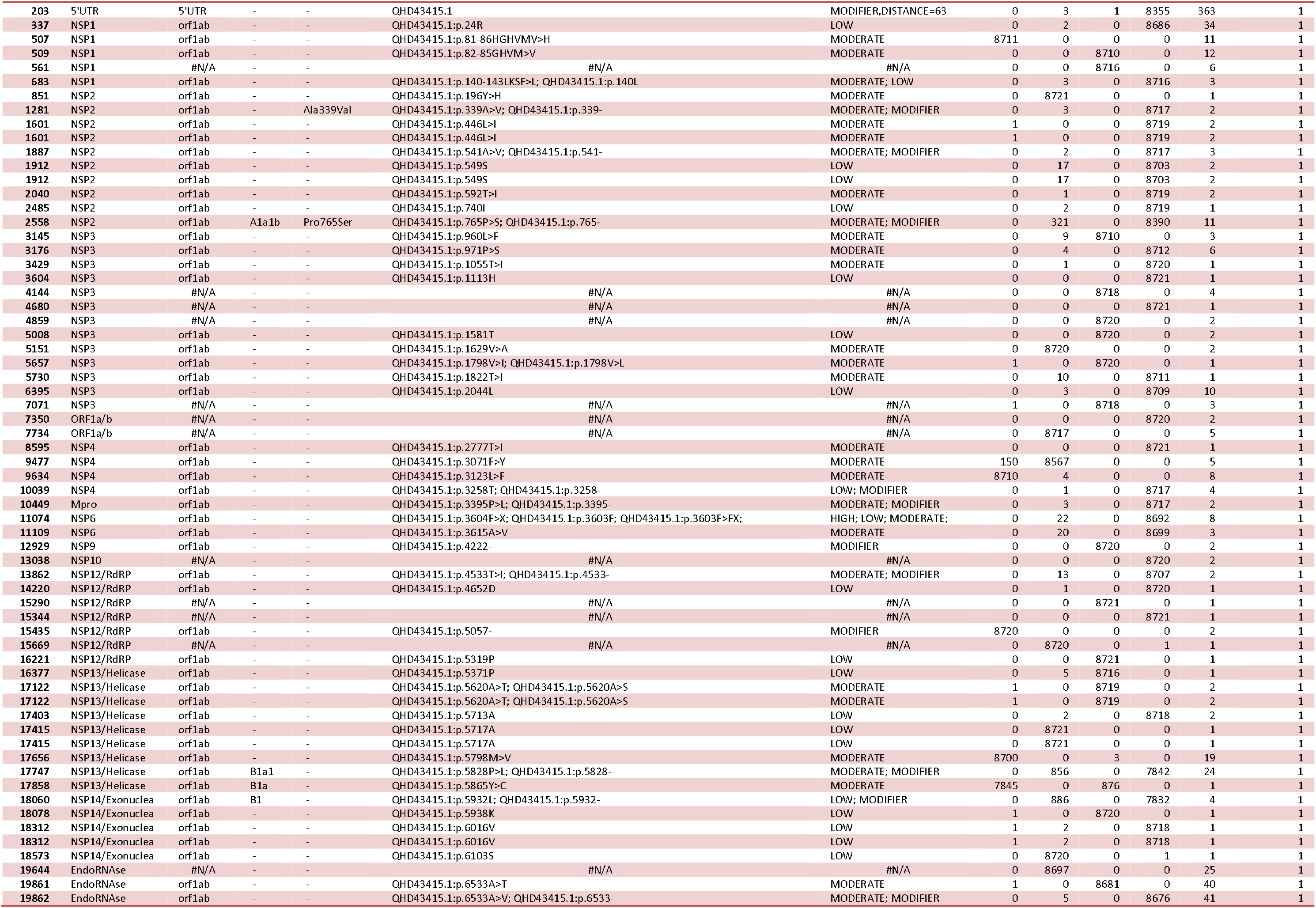
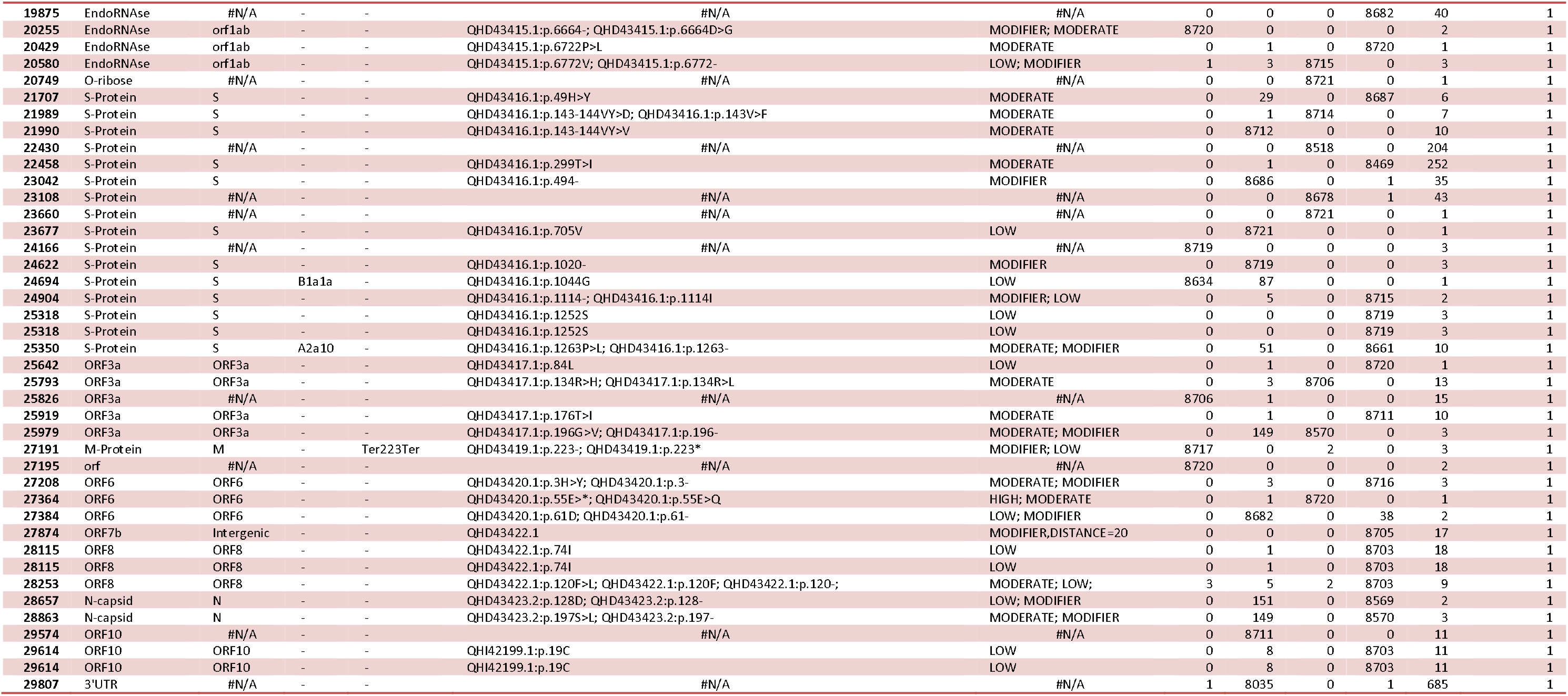
Frequency and description the variations obtained in Cov2 genome from major identified cluster from 104 sequences

### Sample collection and Viral RNA isolation

The nasopharyngeal and oropharyngeal swabs (in viral transport medium) received at NCDC, Delhi through IDSP were subjected to viral inactivation followed by RNA extraction using QIAamp Viral RNA Mini Kit (Qiagen). To ensure that sub-optimal RNA samples are also included in the study, we made use of SuperScript IV (Cat. No. 18091050, Thermo Fisher Scientific, Waltham, MA, USA), for superior first strand cDNA synthesis and included them for sequencing.

### Molecular diagnosis of COVID-19

Reverse transcribed (RT)-PCR assay was used on purified RNA for detection of SARS-CoV-2 in the samples. Quantitative RT-PCR was carried out using Taqman assay chemistry on ABI7500 platform. The primer/probe concentrations and reaction conditions for diagnostics were as per the WHO protocols (Corman et al., 2020). Two target genes were used for diagnosis of SARS-CoV-2, envelope (E) gene for screening and RNA dependent RNA polymerase (RdRp gene) for confirmation. The positive samples were analyzed based on the country of origin (traveler), contact with positive case, geographical location (community), gender and age. Samples from each group were selected and further processed for WGS of the SARS-CoV-2.

### WGS of SARS-CoV-2

#### cDNA synthesis, ONT library preparation and sequencing

cDNA synthesis, ONT library preparation and sequencing has been mentioned in detail in supplementary section. Briefly, 50 ng of total RNA was used for cDNA (first and second strand) synthesis using random hexamer.

A 100ng of double stranded cDNA was used for NGS using a highly multiplexed PCR amplicon approach on the Oxford Nanopore Technologies (ONT, Oxford, United Kingdom) MinION using V3 primer pools (ARTIC Protocol). The amplicons were end repaired, ligated with native barcodes (EXP-NBD104 and EXP-NBD114, ONT) and purified using Ampure beads. The purified product was used for adaptor ligation using Adapter Mix II and purified using a combination of short fragment buffer and Ampure beads. The library was quantified using the Qubit dsDNA HS assay kit and 70ng of the library was used for sequencing. Barcoding, adaptor ligation, and sequencing were performed on samples with Ct values between 16-31. Sequencing flowcell was primed and used for sequencing using MinION Mk1B.

### Illumina library preparation, sequencing and data analysis

The details of methodology for illumina library preparation, sequencing and data analysis are mentioned in supplementary material. In brief, the cDNA pool used for Nanopore sequencing libraries was also used for Illumina library preparation and subsequent sequencing employing Nextera XT protocol. Illumina’s MiSeq platform was used for sequencing. The raw reads were quality checked, trimmed and the mapped human transcripts were removed. The unmapped reads from the human were used to align to the SARS-CoV-2.

### Phylogeny and network analysis

The fasta sequences were aligned using MAFFT considering the MN90894.3 version as the reference sequence. Phylogenetic tree has been constructed using the Neighbour joining algorithm as statistical method and Maximum Composite Likelihood as model in MEGA10 software. FIGTREE was used for the graphical visualisation of Phylogenetic analysis (http://tree.bio.ed.ac.uk/software/figtree). Pheatmap and complex heatmap packages from R were used to plot the heatmaps. The Haplotype Network analysis has been done using PopART [Leigh and Bryant, 2015].

### Protein based annotation and 3-Dimensional protein models

The protein-based annotation was performed according to the reference genome of SARS-CoV-2 (NC_045512 in the NCBI database) to categorize the specific amino acid change. Computational protein structure models of SARS-CoV-2 was created to map high frequency mutations on Spike, Nucleocapsid and nsp3 and RdRp (nsp12) proteins. The detailed methodology for the protein based annotation and 3-dimensional protein models have been mentioned in the supplementary materials.

## RESULTS

### Demographic details of the subjects

The mean (standard deviation) age of the total 127 subjects was 41.4±17.5 years with age range 0.5-76 years and median of 39 years. The gender ratio of male: females in the age group <39 years was 35:28 while the remaining 46 subjects >40 years had the ratio of 58:6.

### Geographical location and travel history

The majority of the SAS-CoV-2 positive samples were obtained from New Delhi covering the national capital region of Delhi, India and few clusters identified as per surveillance team (covering various states of Delhi, Tamil Nadu, Maharashtra, Uttar Pradesh, Andhra Pradesh, West Bengal, Bihar, Orissa, Rajasthan, Haryana, Punjab, Assam and Union territory of Ladakh). The exposure to the COVID-19 was suggestive of travel history of subjects to Europe, West Asia and East Asia. Few subjects were from foreign countries i.e. Indonesia (n=14), Thailand (n=2) and Kyrgyz Republic (n=2). The identified localities of the subjects will further help in molecular surveillance of SARS-CoV-2 in respective geographical regions.

### Profile of SARS-CoV-2 Genome Sequences

#### Amplicon coverage

The average amplicon coverage for the V3 ARTIC primers used in the study was more than 1000X coverage across the majority of the samples (**Figure 1A**). We observed that 73 out of 98 amplicons had more than 1000X in 90% of the samples. Conversely, there were only 25 amplicons which had less than 1000X coverage in 10% of the samples used in the study.

**Figure 1:**
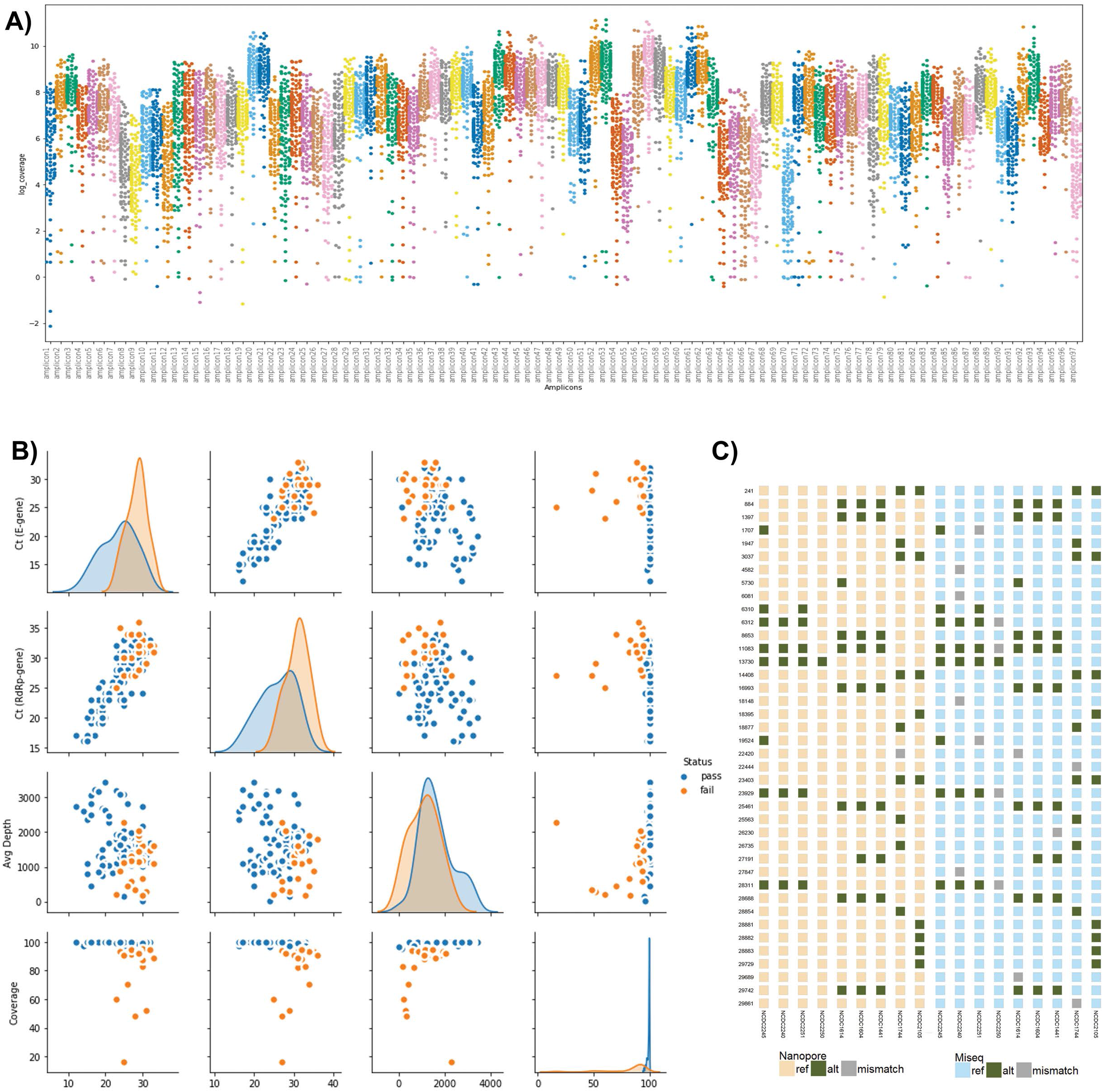
Sequencing data quality parameters and orthogonal platform validation. **A) Arctic Amplicon coverage plot.** It represents the amplicon wise genome coverage of the SARS-CoV-2 genome. 73 out of 98 (~71.5%) amplicons have more than 100X coverage in 90% of the sequenced samples. **B) Genome coverage and sequencing depth of the SARS-CoV-2 genome.** We plotted the Ct value of the two genes (E-gene and RdRp gene) used for RT-PCR vis-a-vis genome %age coverage and sequencing depth of the samples (Blue - missing N% below 5 and Orange - N% above 5). In general, it was observed that samples with lower Ct value have higher average sequencing depth and genome coverage compared to higher Ct value samples. **C) Genetic variants across ONT and Illumina platform**. A subset of ONT sequenced samples were sequenced on MiSeq to benchmark the genetic variants observed in the positive samples. The genetic variants marked in yellow are the same between the two platforms.

#### Genome coverage vs Ct value

We also looked into whether lower Ct values are a good indicator of genome coverage using a minimal set of virus mapping reads. We plotted genome coverage and average sequencing depth across Ct value of both the genes (E and RdRp). It was observed that higher Ct values (27 onwards) have increased possibility of lower genome coverage (**Figure 1B**) although some lower Ct value samples also had incomplete genome coverage. This may be due to the integrity of the RNA used for sequencing. It was observed that the average sequencing depth was higher (1000X-3000X) for samples with lower Ct values, whereas higher Ct value samples have relative lower average sequencing depth (100X-1000X). This would be important for informed decision making for reads required for ONT based sequencing.

#### Sequencing platform comparison

We sequenced a subset of samples on orthogonal platforms and sequencing methods (shotgun and amplicon) using ONT and Illumina platform. Significantly, we observed that the genetic variants were common between both the platforms (**Figure 1C**).

### NGS analysis for SARS-CoV-2 sequence

A total of 104 samples passed the quality threshold for mapping full genome coverage threshold for SARS-CoV-2 genome (accession ID:) <0.05 N content with median coverage-~1500X.

Twenty-three samples that did not qualify the threshold criteria were excluded from strain identification. However, few samples were retained for variant analysis wherever the sequencing coverage was sufficiently of high quality for variant calling.

### Phylogeny and variant analysis

The phylogenetic analysis of 104 high quality sequences reveal all the strains to be grouped into two major clades, a sub-clade and other clades (**Figure 2** and **Supplementary Figure 2**). In total we observed 163 variants representing singletons (107 variants), rare: 2-5% (45 variants), and common variants: >5% (11 variants). The common variants observed were 241 (Leader sequence), 3037 (NSP3), 6310 (NSP3), 6312 (NSP3), 11083 (NSP6), 13730 (NSP12/RdRp), 14408 (NSP12/RdRp), 23403 (S-Protein), 23929 (S-Protein), 25563 (ORF3a), 28311 (N-capsid).

**Figure 2:**
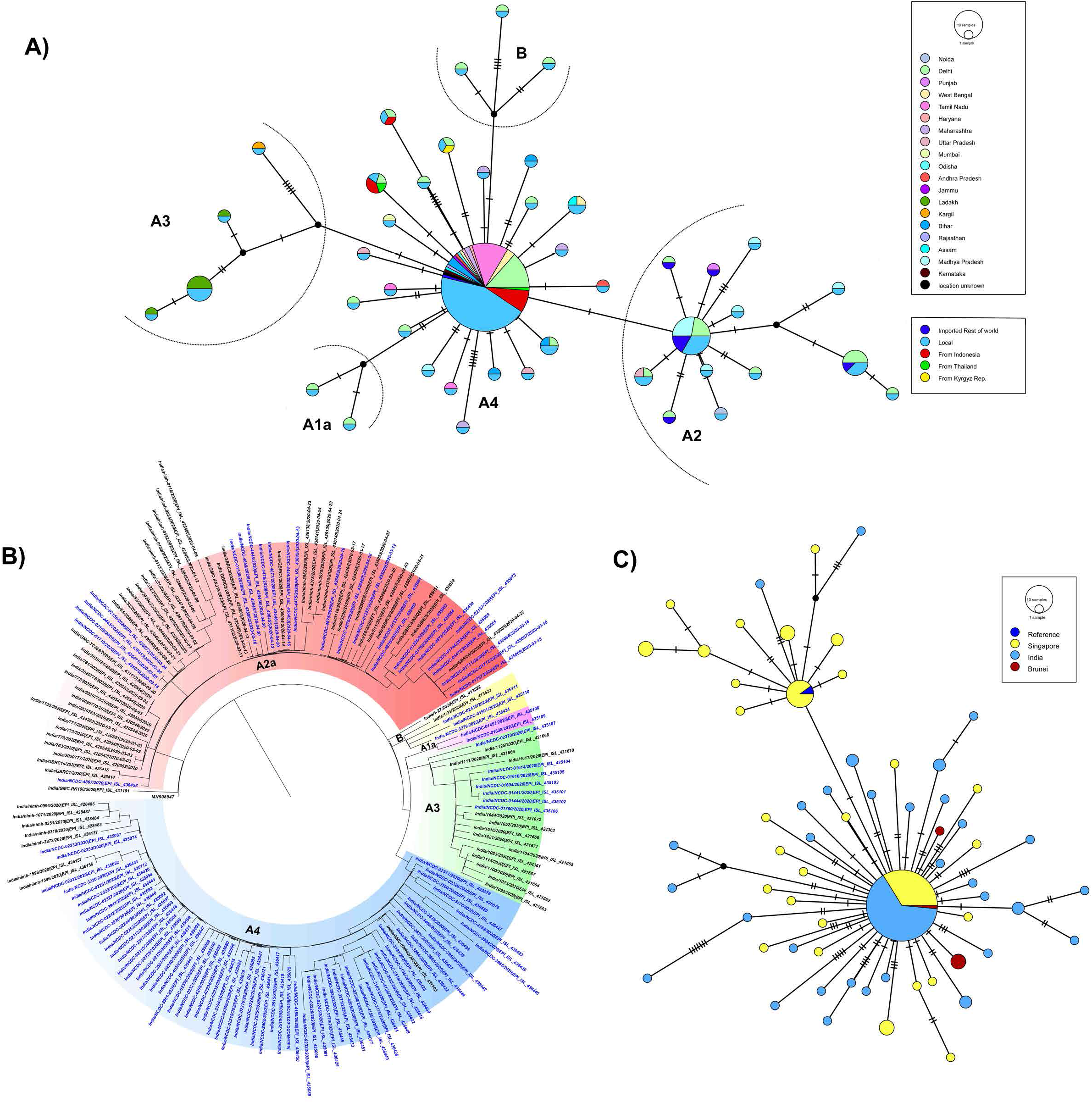
Haplotype network and Phylogenetic analysis of SARS-CoV-2 sequences. **A)** The network analysis of SARS-CoV-2 sequences from this study showing distinct clades with their geographical locations. A4 clade described in this study for the first time has widespread geographical affiliations. A3 being more confined to Ladakh and mostly these cluster represents introduction of the infection through travel history. **B)** phylogenetic tree of SARS-CoV-2 genomes from sequences submitted from all across India depicting major clade based distribution of SARS-CoV-2 in India. **C)** Network analysis of A4 clade sequences submitted from Indian and neighbouring East Asians countries.

#### Cluster-1

This cluster of these sequences (n=26) have shown G-clade [(Global Initiative on Sharing All Influenza Data, GISAID based nomenclature for variant at 23403 (Spike protein: D614G) position]. Based on Nextstrain classification 26 sequence belonged to A2a clade [(denoted by positions C241T; C3037T, A23403G (S: D614G); C14408T (ORF1b/RdRp: P314L)] (**Figure 2A**). The additional frequent variants in this cluster observed were 25563 (ORF3a, n=9),18877 (NSP14/Exonuclease, n=7), 26735 (M-protein, n=6), 22444 (S-protein, n=5), and 28854 (N-capsid, n=5). One novel variant *1947 T>C (NSP2)* was observed in two strains in this cluster.

#### Cluster-2

In our study large numbers of strains (n=65) belong to this unclassified cluster (as per GISAID and Nextstrain). The strains in this cluster had the predominant variants, G11083T variant (NSP6) (n=65), C13730T in RdRp (n=65), C28311T (N-capsid); n=65, C6312A (NSP3 variant); n=64 (one sequence being called N), C23929T (S-protein), n=50 (other being low depth/N bases). The variant C6310A (NSP3); n=22 being observed as another frequent alteration (**Figure 2B**). We also observed few novel variants, *G12685T(NSP8)*; n=5 and *TC1706T(NSP2)*; n=4, *T7621C (ORF1a/b)*; n=3, *A21792T (S protein)*; n=3, *G13920A (NSP12/RdRp)*; n=2, *A16355G (NSP13/Helicase)*; n=2 and *G18803T (NSP14/Exonuclease)*; n=2 in this cluster.

The majority of the key cluster variants 11083, 13730, 28311, 6312, 23929 are also shared in sequences submitted from Singapore region and Brunei (GISAID) and also similar clade sequences were observed in India submitted by National Institute of Mental Health and Neuro-Sciences (NIMHANS) and Gujarat Biotechnology Research Centre (GBRC) cohort. From history, based on the geographical location of the subjects of this cluster, a significant number of natives of Indonesia (n=7) and two each from Thailand and Kyrgyzstan were part of this cluster from our study site. This probably suggests introduction of this particularly from East Asian countries into India.

#### Cluster-3

This subclass of strains (n=7) harboring a common variant G11083T (NSP6), G1397A (NSP2) and T28668C (N-capsid) are described for the A3 clade (Nextstrain) in additions to G29742T (**Figure 2A**). Other mutated positions i.e. C884T (NSP2), G8653T (NSP4) were observed in 5 samples, whereas T16993C (NSP13), n=4; T25461C (ORF3a), n=4 and A27191G (M-protein), n=2 are putative novel sites.

The phylogeny analysis of these clusters segregated with the other Indian SARS-CoV-2 genome sequences as recently reported (GISAID) (**Figure 2B**).

#### Other SARS-CoV-2 genomes

Two SARS-CoV-2 belonging to A1a clade had a SNP profile of 11083(NSP6)/14805 (NsP12/RdRp)/2480 (NSP2)/2558 (NSP2)/26144 (ORF3a). In addition, we observed three B clade sequences having position 8782 C>T (NSP4) and 28144 T>C (ORF8; S clade GISAID) mutated and with one sequence with an additional C18060T B1 variant. One genome from Maharashtra had no variants and probably represented the first genome sequenced from Wuhan, China.

### Re-defining Cluster-2 with neighbourhood re-joining

With over represented variants in cluster-2 for variants 11083/13730/28311/6312/23929, we defined this cluster with A4 clade. This has similarity with sequences submitted from Singapore, Brunei and other Indian sequences submitted. The haplotype network analysis suggests that these sequences are having a common origin from East Asia/South-East Asia (**Figure 2C** and **Figure 3**). This A4 clade has multiple variants in important region of viral genome, RdRp (A97V), N-capsid (P12L), NSP3 (T2016K), NSP6 (L37F) and NSP3 (S1197R) variants. In our cohort of samples, the majority of subjects were from Tamil Nadu, Delhi and Indonesia and others were from various other states (**Figure 3**).

**Figure 3:**
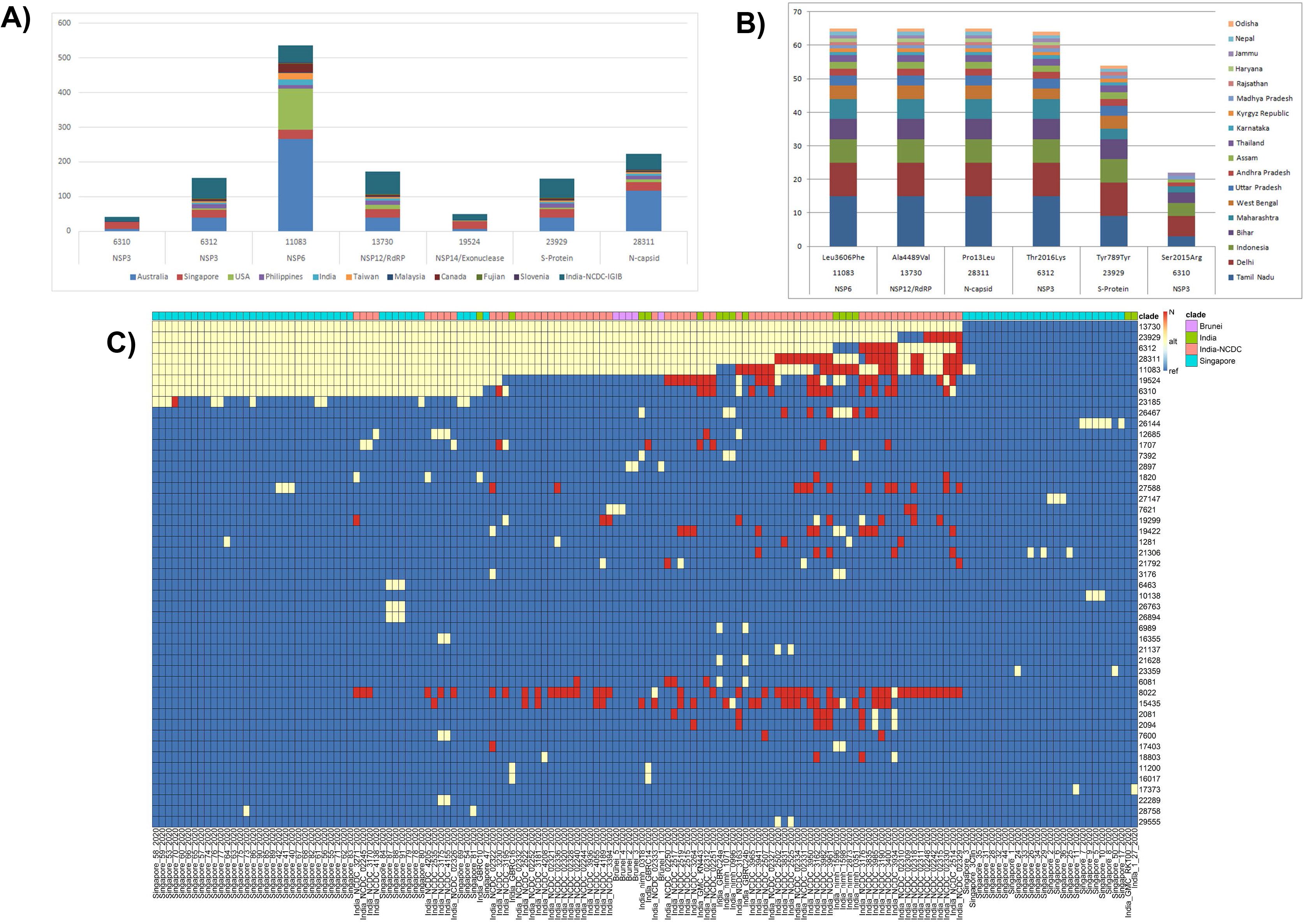
Distribution of A4 clade within India and globally. **A**) Distribution of A4 clade variants across different geographical regions from the cohort. **B)** Distribution of A4 clade variants across different geographical regions across the globe. **C) C**omparison of A4 cluster sequences across the South-East Asian region showing sharing of variants and haplotype across South-East Asian region.

### Protein-wise analysis of SARS-CoV-2 variants

To provide quantitative insights into the mutant proteins, we characterized amino acid substitutions across the 104 viral genomes. Of the 53 point mutations identified, 29 were missense that resulted in amino acid substitutions. **Figure 4** plots the occurrence of these mutations as a function of each viral protein. The frequency of amino acid variations were maximum in nsp6 (L37F) present in 68 genomes, followed by nsp12 (A97V) in 65, nsp3 (T1198K) in 62 and nucleocapsid (P13L) in 53 genomes. Interestingly, D614G mutation in spike protein which is considered as a prevalent global mutation [Korber et al., 2020; Chandrika et al., 2020], was present in only 26 of the 104 sequenced genomes.

**Figure 4:**
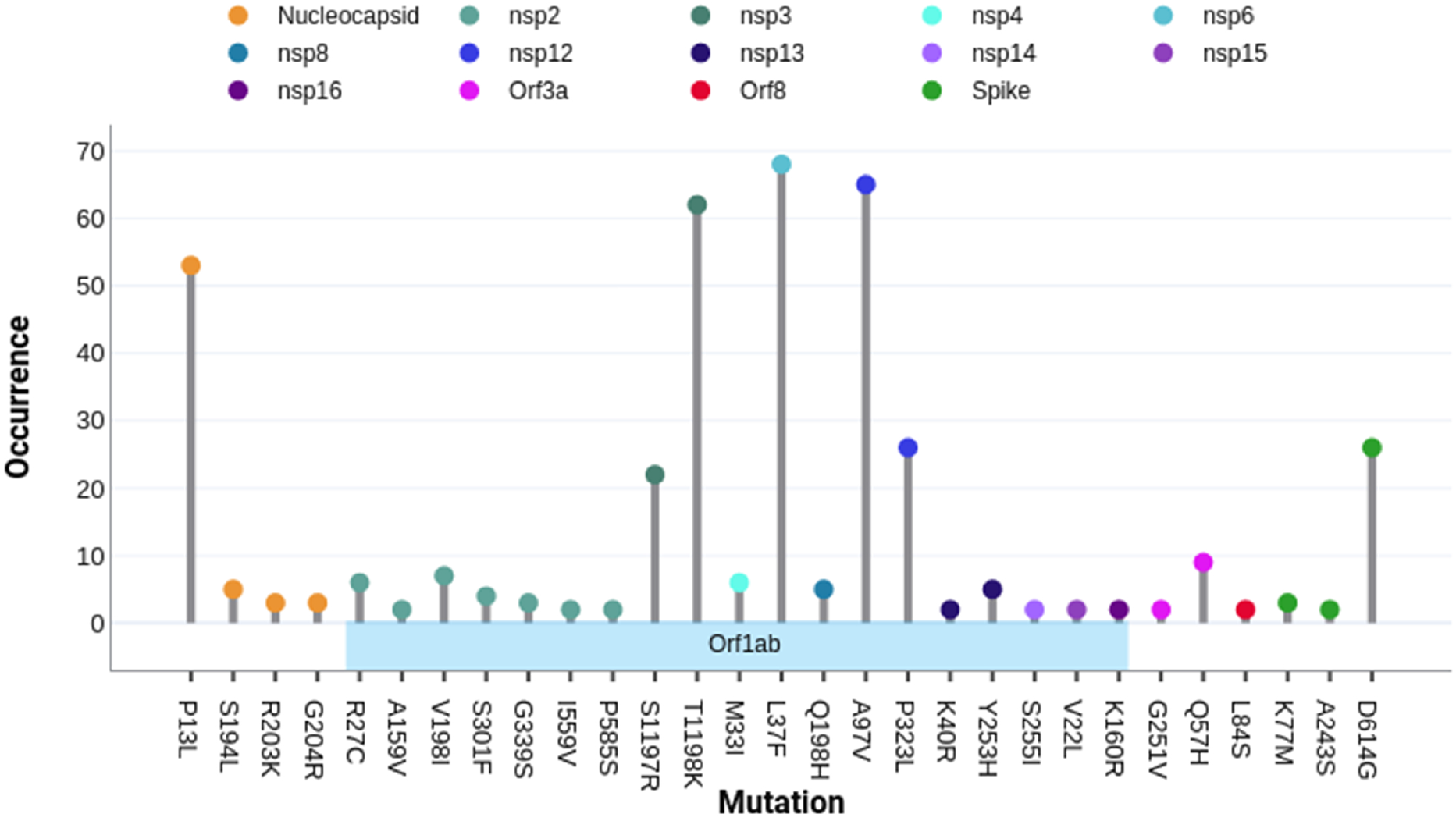
Protein annotation of the amino acid substitutions. 29 Amino acid substitutions obtained from 104 sequenced genomes were mapped on various viral proteins as a function of frequency of these missense mutations. The majority of mutations are in Orf1ab region, which is decomposed into multiple non-structural proteins. The color mark at the end of occurrence bar indicates the protein name shown in the above legend.

Next, we correlated the occurrence of each mutation with the type of amino acid change (**Figure 5A**). Interestingly, ~45% of these mutations showed no change in the nature of the amino acids, suggesting that the viral proteins harbors subtle changes in the protein shape or function. Within high-occurring mutations, P13L, L37F, A97V also showed no major residue alterations. However, there are other loci that involve complete change in amino acid type once mutated such as interchange between polar, hydrophobic or charged. For instance, high frequency positions, including T1198K in nsp3 involve acquisition of a charged group. In addition, the key mutation in spike protein (D614G) also involves loss of the charged group. These mutations that lead to positively charged groups may cause more severe structural and functional effects.

**Figure 5:**
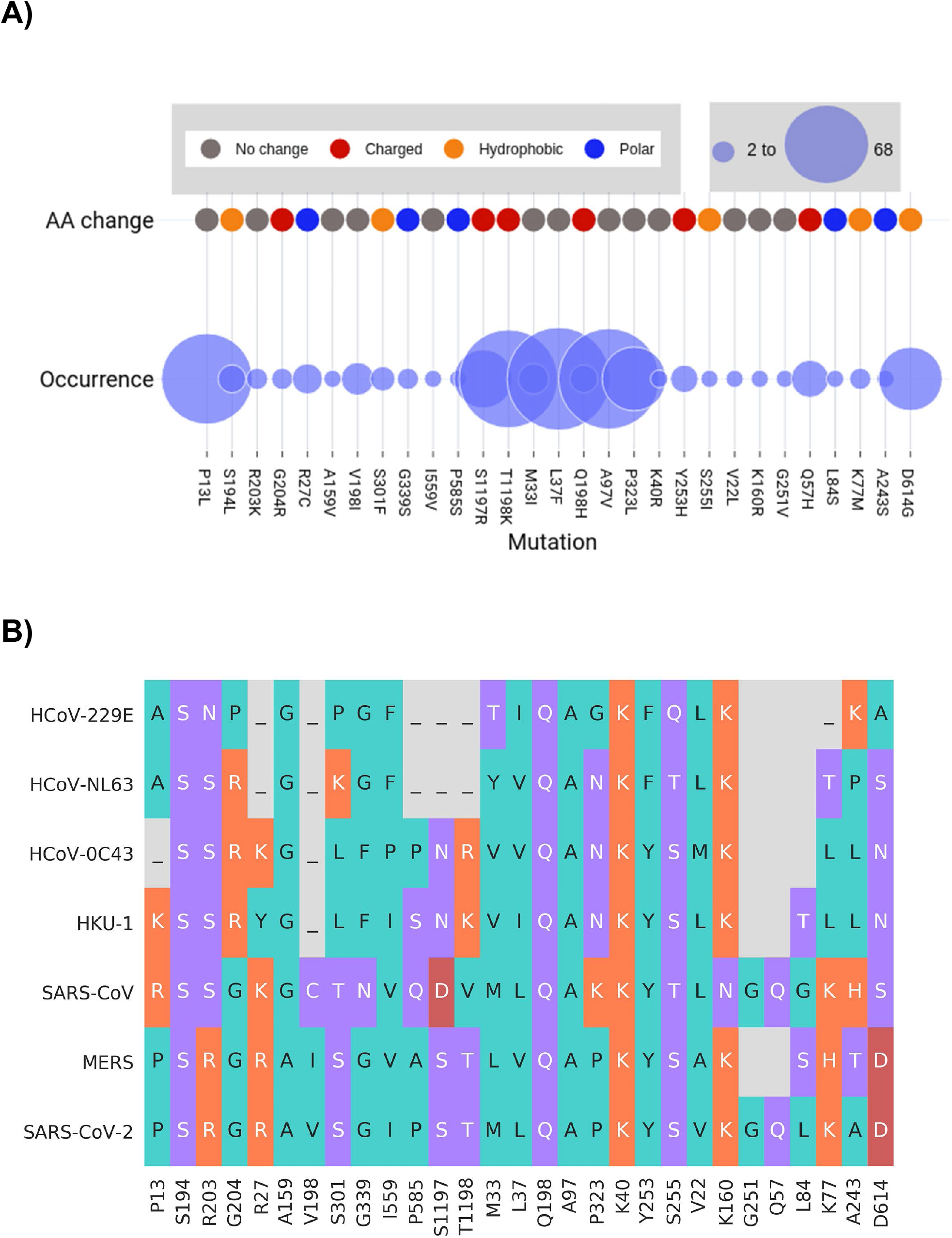
Amino acid properties marked as a function of mutations. The first row shows amino acid change, with polar, hydrophobic, charged, and unchanged residues shown in violet, blue, green, and yellow, respectively. The second row shows the occurrence of protein mutations in 104 genomes, with the larger circle representing higher occurrence (69) and the smallest circle shows the presence in 2 genomes. The last row depicts the conservation score of that particular mutation of SARS-CoV-2 with 6 other coronavirus. The conservation ranges between 1 to 9 with 9 depicting the highest score.

We also compared SARS-CoV-2 mutation sites with other six coronavirus sequences (**Figure 5B**). Most of the mutations were present mostly in variable locations. Out of 29 mutations, 10 are present on highly conserved residue locations. Interestingly, higher frequency mutations are at positions that evolve faster/are variable across the coronaviruses except A97L and L37F, which are present on conserved locations.

To understand how these mutations are changing the local environment, we mapped higher frequency mutations onto SARS-CoV-2 protein structures, including nucleocapsid, nsp3, nsp12, and spike protein (**Figure 6**). In nucleocapsid, there are two structural domains, the N- and C-terminal connected via a long linker region that facilitates RNA packing [Kang et al., 2020]. We found that all the mutations, including P13L found in 53 genomes, are present outside these domains, specifically, linker regions (S194L, G204R, R203K), and the N-arm of the protein (P13L). The nsp12 is a highly conserved protein with multiple domains. The observed mutations are overlaying onto the interface (P323L) and NiRAN (A97L) region. The latter is critical as it contains a Zn+ binding site, however, little is known about the exact functional output. In contrast, the P323L mutation is present on protein interaction junctions where a hydrophobic cleft is known to bind to inhibitors (**Figure 6B**). This variant will result in loss of kink due to amino acid substitution from proline to leucine, i.e., 5-membered amino acid which resides in the buried area of the protein from to an acyclic amino acid. The nsp3 protein has a similar higher order domain arrangement, and the mutation is present on the NAB domain, which is a nucleic acid binding domain and also interacts with nsp12. This mutation may impact RNA synthesis machinery; however, little is known about its exact mechanism of action. Lastly, the D614G mutation in spike protein is an interesting substitution and has been reported with increased tally [Korber et al., 2020; Chnadrika et al., 2020]. Structurally, this mutation is located in the S1 subunit that also contains the RBD domain. Although present outside the functional region, the proximity of D614G around S1 cleavage site implicates an important change in the local environment.

**Figure 6:**
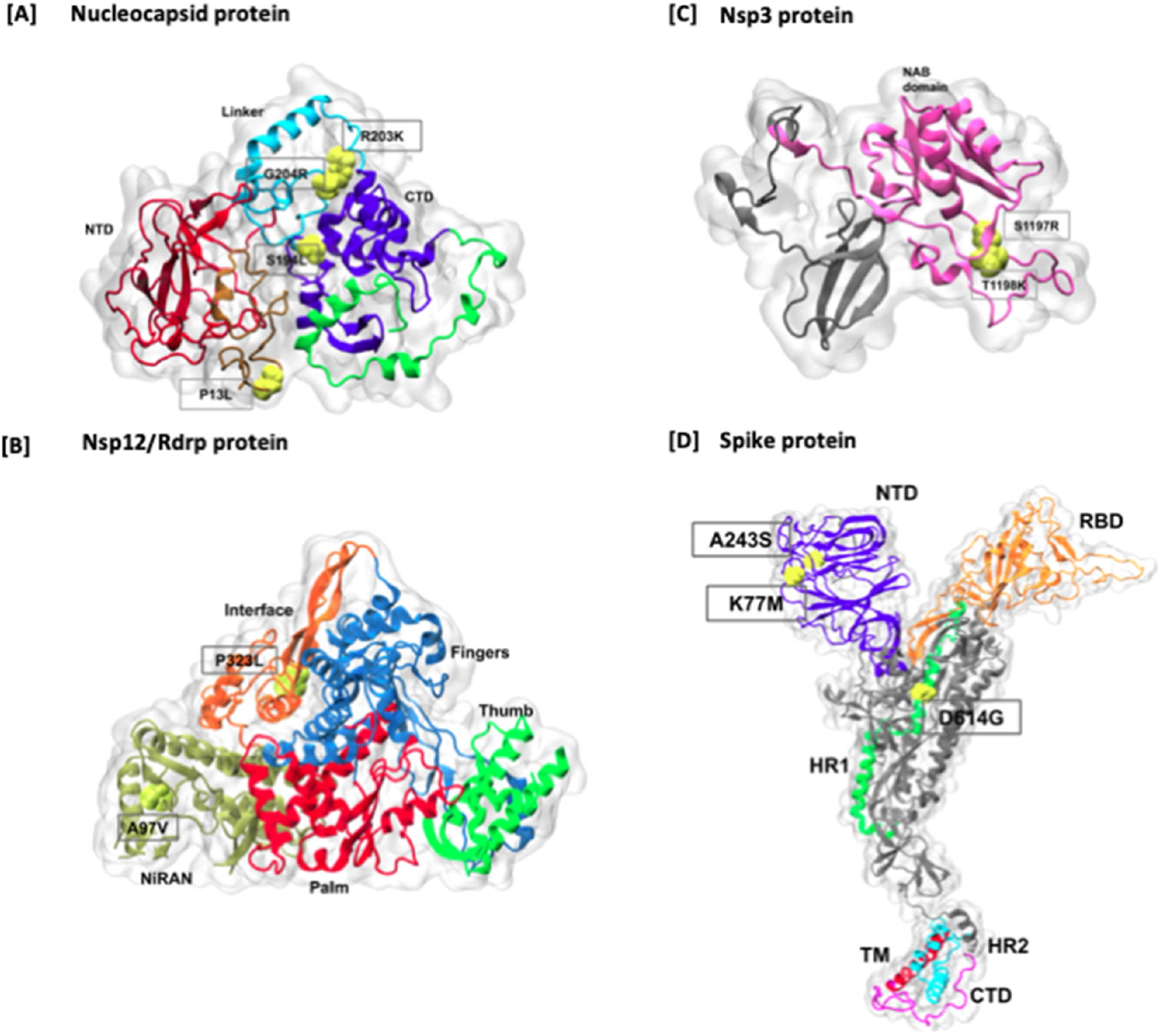
Mapping of higher frequency amino acid mutations on Nucelocapsid and Spike proteins. The mutations are marked in red color on the surface representation of each protein. In Spike protein, all the domains are highlighted in different colors, including NTD, RBD, HR1, Fusion peptide region, HR2, TM, and CT domains. In addition, cleavage sites are also marked onto the structure.

### Geographical distribution of SARS-CoV-2 in India

SARS-CoV-2 classified for their preponderance of distribution across geographical locations of India vis-a-vis states of India. Enrichment and depletion of specific clades/classes was observed in certain states of India. We explored the possibility of whether travel history based viral strain showed signs of adding new variants once it diversified in India.

## DISCUSSION

This is the first comprehensive genomic picture of the SARS-CoV-2 prevalent in the Indian population during the early phase of outbreaks. The understanding is important keeping in view the vast geographical expanse and population density of India. There were three major waves of viral entry in India associated with multiple outbreaks (**Figure 7**). First wave includes importation of SARS-CoV-2 (A2a cluster) through travelers from Europe (Italy, UK, France etc) and the USA. Second wave of SARS-CoV-2 (A3 cluster) was linked with the Middle East (Iran and Iraq). Third wave comprises combined viral (haplotype redefined as A4) entries from Southeast Asia (Indonesia, Thailand and Malaysia) and Central Asia (Kyrgyzstan). The study taken together with other reported genomes (Potdar et al., 2020) revealed A4 cluster (previously unclassified) is the most prevalent in Indian population. Many novel mutations identified may be specific to Indian conditions but more genomic data is needed to strengthen the assumption to rule out sampling bias and other factors (Lu et al., 2020). During the early phase of the epidemic in India, imported cases followed by local transmission were restricted to urban regions where effective IDSP network, diagnostic support, health care services and timely placed interventions (nationwide lockdown, establishment of containment zones and practicing social distance) interrupted community transfer. The mounting burden of epidemic in urban regions associated with job crunch forced migrant workers to return to their home mostly situated in rural areas which is an impending threat of the viral entries in rural regions (comprising major portion of the Indian population) and expected community transmission. To tackle domestic transmission at larger scale and higher risk of community transfer, preventive measures shall be strengthened in rural regions in addition to safe transportation arrangements for domestic migrants.

**Figure 7:**
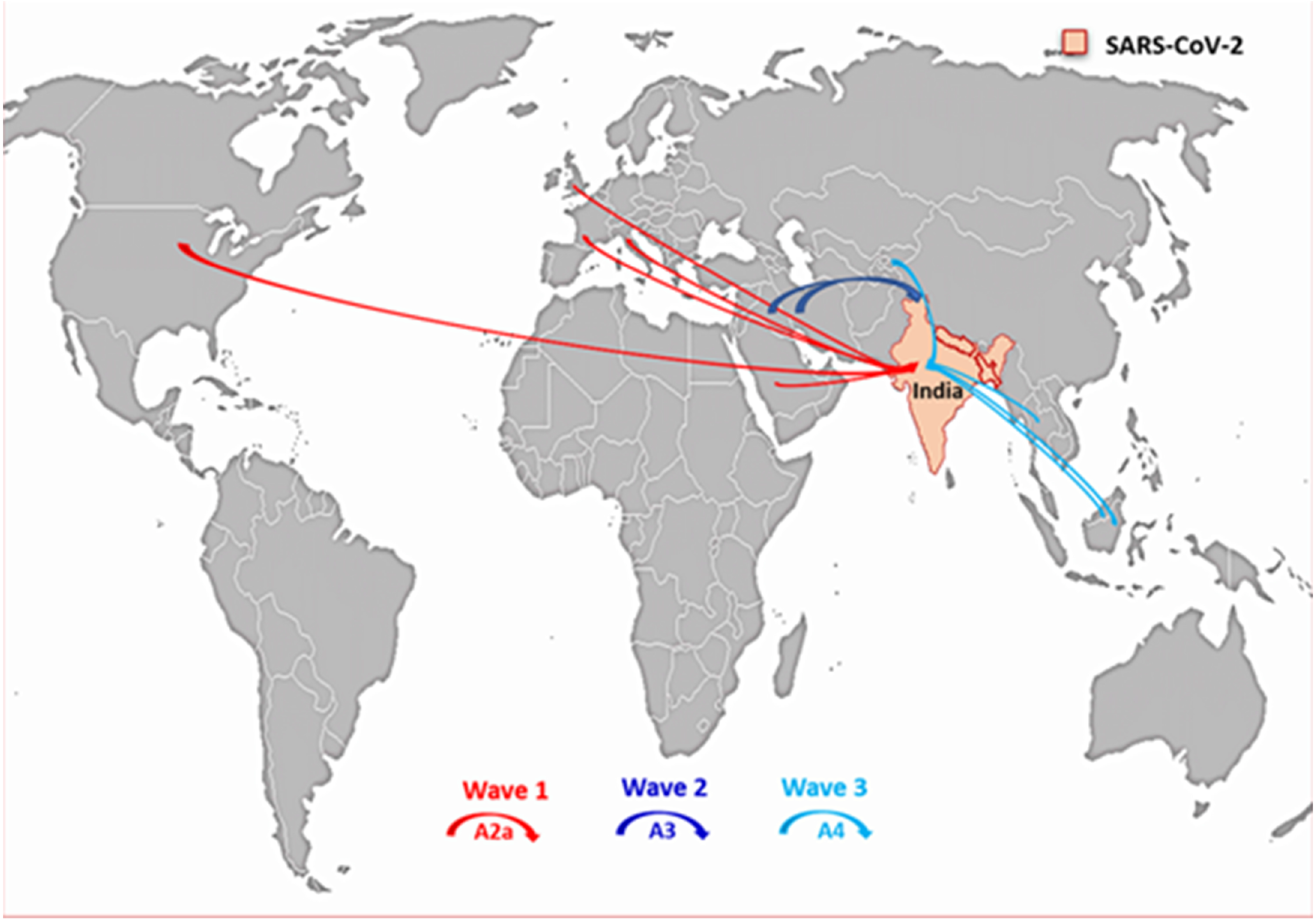
Scheme showing importation of prevalent SARS-CoV-2 genomes (3 major waves) in India.

The national lockdown may have led to evolution/gain of certain variants with potential role in adapting to Indian conditions. This may have given rise to a distinct lineage awaiting its inclusion in Nextstrain (an open-source project to harness the scientific and public health potential of pathogen genome data). A more detailed analysis of these genomes might provide information whether these variations need to be considered during design of diagnostic primers as the need for testing shoots up. It may allow for creation of cost effective panels to trace the movement of lineage specific strains across geographical regions more rapidly and effectively. Lots of efforts are ongoing to identify suitable vaccine candidates through docking studies. These observations are important to consider the variants especially that map to the Indian genomes during such prioritization studies since these strains would now form a major fraction of the genomes that are likely to become more prevalent in India after lockdown. Mapping of these variant genomes in conjunction with the clinical history in terms of recovery, hospitalization and co-morbidity might allow identification of variants that should be actionable and would also have relevance for prognosis. It is imperative that robust genomic data based on large sample size including rural populations with even distribution can bring out the real scenario once correlated with epidemiological data eventually helping in drafting of further management policies.

## SUPPLEMETARY INFORMATION

Supplementary Material, Supplementary figures 1: location map; supplementary figure 2: heat map representation of SARS-CoV-2 variants per sample in respective clusters.

## ACKNOWLEDGEMENTS

NCDC greatly acknowledge the support of Prof. Dr. Christian Drosten, Charite – Universitatsmedizin, Berlin for promptly providing the positive controls for qPCR. The authors do acknowledge GISAID for sharing the genomic sequences in public domain and other contributors SARS-CoV-2 genomic data. We would like to gratefully acknowledge the efforts of officials involved in IDSP network and associated hospitals in sample collection and timely data sharing. We would like to thank the financial aid provided by Ministry of Health and Family Welfare, Government of India. CSIR-IGIB would like to acknowledge Genotypic Technologies Pvt. Ltd., Bangalore, India for its role in facilitating ONT sequencing. Authors also acknowledge the role of all the technical and support staff of NCDC involved in COVID-19 testing and Subhash Gurjar (CSIR-IGIB) for facilitating reagents procurement amidst lockdown and for providing other assistance in the laboratory work. We thank CSIR, India for funding support towards SARS-CoV-2 genome sequencing.

## AUTHOR CONTRIBUTIONS

A.A., P.K., P.R. and S.K. conceived the study; P.K., M.S.D., M.B. and P.R. planned and optimized diagnosis; M.S.D., H.V., U.S., Priyanka S., M.D., P.G., D.K., S.B. performed viral inactivation and diagnosis; S.S., H.L., A.T., B.N., V.J., H.G., P.M., N.S. and Sujeet S., did data compilation, field work and epidemiological analyses; R.P. and M.F. planned the sequencing experiment; P.S., S.W., N.T., R.P. and M.F. conducted the sequencing experiments; V.A., B.U. and M.F. analysed the sequencing data, Prateek S. and D.D. did independent sequencing analysis validation, S.F., D, S.R., N.J. and L.T. performed the protein analysis; P.K., M.F. and R.P. interpreted final data and wrote the manuscript; M.M., M.S.D. and M.B. edited the manuscript. All the authors read and approved the final manuscript.

## DECLARATION OF INTERESTS

Authors declare no conflict of interests with the present study.

## Notes

### Competing Interest Statement

The authors have declared no competing interest.

