## Supplementary material, supplementary figure 1, supplementary figure 2 for "Integrated genomic view of SARS-CoV-2 in India": Supplementary file_bioRxiv.docx

**Supplementary Materials**

**cDNA synthesis:** Total RNA from SARS-CoV-2 positive samples were quantified using Nanodrop and 50ng of the RNA was taken for double-stranded cDNA synthesis. First strand cDNA was made using 1.0μl of random hexamer (50ng/μl), 1.0μl of dNTPs (10nM) and 13.0μl of total RNA with volume adjusted with nuclease free water (NFW), followed by incubation at 65ºC for 5 mins and cooling on ice. To this, 4.0μl of 5X SSRT IV Buffer, 1.0μl of 100mM DTT, 1.0μl of Ribonuclease Inhibitor and 1.0μl of SSRT IV Enzyme (200U/μl) was added (Cat. No. 18091050, Thermo Fisher Scientific, Waltham, MA, USA) with incubation at 23ºC for 10 minutes, 50ºC for 10 minutes and 80ºC for 10 minutes. 1.0μl of RNase H was added to this and incubated at 37ºC for 20 mins. 20.0μl of first strand cDNA was heated at 95ºC for 3 minutes after addition of 10pmol of random primers, 10μM dNTPs and 1X Klenow Buffer, followed by immediate cooling on ice. Soon after, 1.0μl of Klenow Fragment (Cat. No. M0210S, New England Biolabs) was added with incubation at 37ºC for 60 mins, 75ºC for 10 mins and 4ºC for 10 mins. This was followed by Ampure beads purification (Cat. No. A63881, Beckman Coulter) and quantification using Qubit dsDNA HS assay kit (Cat. No. Q32854, Invitrogen).

**ONT library preparation and sequencing:** 100ng of double stranded cDNA was taken for Next Generation Sequencing (NGS) using a highly multiplexed PCR amplicon approach for sequencing on the Oxford Nanopore Technologies (ONT, Oxford, United Kingdom) MinION using V3 primer pools (ARTIC Protocol). Amplification was done using TaKaRa LA Taq® DNA Polymerase Hot-Start Version (Cat. No. RR042B, Takara) along with 2.5mM dNTPs, 10X Buffer II LA Takara, 10μM primer pool with final volume adjusted to 25μl using NFW. Primers were made into two pools - pool A and pool B, with 5μl of each primer from the 100μM primer stock. The stock (100μM) was diluted with NFW in order to obtain a working stock of each pool at 10μM. PCR was performed with initial denaturation at 98ºC for 30 secs followed by denaturation at 98ºC for 15 secs, 65ºC for 5 mins, for a total of 35 cycles, with hold at 4ºC. Post PCR pooling (pool A and pool B) of PCR amplicons was followed by purification using Ampure beads. Post purification, 1.0μl of the library was run on DNA1000 Agilent bioanalyzer (Cat. No. 5067-1504, Agilent) to check for a size of ~400bps. 125ng of each sample was taken forward for End prep with NEBNext Ultra II End Repair/dA tailing module (Cat. No. E7546, New England Biolabs). The reaction mix was incubated on a thermal cycler at 20ºC for 5 mins followed by 65ºC for 5 mins. 1.5μl of End prep DNA was taken forward for Native Barcode Ligation using native barcodes (EXP-NBD104 and EXP-NBD114, ONT) and Blunt/TA Ligase master mix (Cat. No. M0367, New England Biolabs). The mix was incubated at room temperature for 15 mins followed by purification using Ampure beads. The purified product was used for adaptor ligation using Adapter Mix II and Quick T4 DNA ligase (Cat. No. M2200L, New England Biolabs). Post adaptor ligation, it was purified using a combination of short fragment buffer and Ampure beads resulting in a sequencing ready library. Library was quantified using the Qubit dsDNA HS assay kit (Cat. No. Q32854, Invitrogen) and 70ng of the library was used for sequencing. Barcoding, adaptor ligation, and sequencing were performed on samples with Ct values between 16-31. A ‘no template control’ was created at the cDNA synthesis step and amplicon generation step to detect cross-contamination between samples. Controls were barcoded and sequenced with both the high and low titer sample groups. Sequencing flowcell was primed and used for sequencing using MinION Mk1B.

**Illumina library preparation and sequencing**

Common pool of cDNA was used for making both Illumina and Nanopore sequencing libraries and subsequent sequencing. 100ng of cDNA was used for making Illumina library using Nextera XT protocol. Tagmentation of cDNA was done which tagged and fragmented the cDNA by addition of Amplicon Tagment Mix (ATM) and tagment DNA buffer with incubation at 55ºC for 5 mins with heated lid option. Tagmentation was stopped by addition of neutralization tagment buffer. This was followed by the addition of unique index adapters (i7 and i5 adapters) to the samples. Index adapters are then used for PCR amplification at 72ºC for 3 mins, 95ºC for 30 secs and 12-cycles of 95ºC for 10 secs, 55ºC for 30 secs, 72ºC for 30 secs; and 72ºC for 5 mins. The PCR product was purified using AgencourtAMPure XP beads. The quantity of the sequencing ready library was measured using Qubit dsDNA HS assay kit (Cat. No. Q32854, Invitrogen) and quality by Agilent DNA HS kit (Cat. No. 5067-4626, Agilent). Illumina’s MiSeq platform was used for sequencing.

**Miseq data analysis**

The raw reads from the miseq has been quality checked by FASTQC. Trimgalore was used to trim the reads containing bad quality and the minimum length of 40 base pairs was kept as a threshold for the reads. HISAT2 is used to map the reads to the human genome (GRCh37) to remove potential the human rRNA reads for the contamination with default parameters [Kim et al. 2019]. The unmapped reads from the human are converted from bam to fastq using bam2fastq to align to the SARS-CoV-2. HISAT2 was used to align unmapped reads with the SARS-CoV-2 reference genome (MN908947.3 build). Using both samtools and bcftools from the SARS-CoV-2 aligned bam files the consensus fasta was generated. The variants in the samples were called using bcftools and varscan.

**Protein based annotation**

In order to categorize the specific amino acid change and the proteins containing the variants, they were annotated by SnpEff version 4.5 [Cingolani et al., 2012]. The annotation was performed according to the known reference genome of SARS-CoV-2 i.e. NC_045512 in the NCBI database [Wang et al., 2020]. SARS-CoV-2 polypeptide ORF1ab encodes 16 non-structural proteins (nsp) as a result of proteolytic processing. Hence, for better mapping of the variants present in ORF1ab, we annotated the variants according to the respective nsp residue number.

Further, conservation analysis of the full-length sequences of proteins harboring these mutations was done on the basis of the six other coronaviruses. The multiple sequence alignment of seven protein sequences was performed by Clustal-Omega [Maderia et al., 2019]. The conservation score of ORF3a and ORF8 were calculated with low confidence due to introduced gaps at these positions during alignment. The amino acid type was defined as hydrophobic (G,A,V,L,I,M,P,F,W), polar (S,T,N,Q) and charged (H,K,R,D,E). With this definition, the type of change of residues was calculated.

**3-Dimensional protein models**

To map the high frequency mutations on proteins, we took computational protein structure models of SARS-CoV-2. Spike protein exists as a homotrimer consisting of 1273 residues in each chain with a total of 3819 amino acids. Electron microscopy structures are available for different conformations of Spike protein [Wrapp et al., 2020; Walls et al., 2020;]. However, these structures contain many missing residues and a major portion of the S2 domain. Similarly, nsp3 also known as PL-PRO (papain like proteinase) is a large multi-domain transmembrane protein of 1945 residues long. Therefore, the complete structural model for Spike, Nucleocapsid and nsp3 were taken from SSGCID models provided by Bakers lab. The models were generated through comparative modelling and by using Robetta protein prediction server [Kim et al., 2004]. For nsp12, we took PDB structure 6M71 where the first 30 residues were missing [Gao et al., 2020]. For mapping the nsp3 mutations, we considered the model for only one domain called NAB i.e. Nucleic acid binding domain (residue 1088-1201) which is conserved in Betacoronavirus [Angelini et al., 2013].

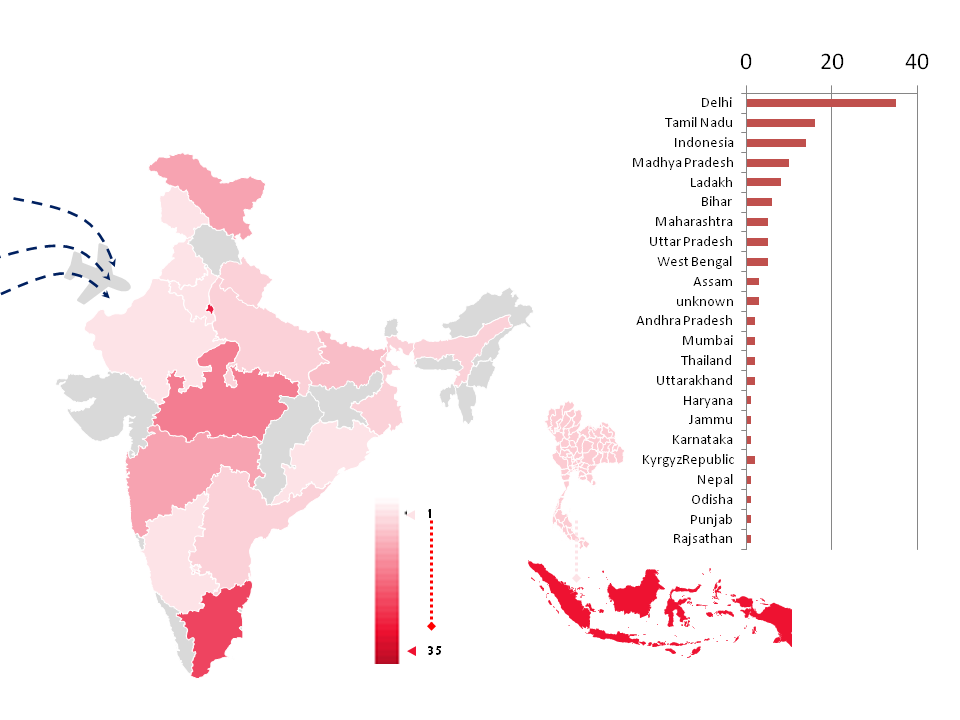

Figure S1: A schematic diagram showing numbers of samples with their geographical affiliations with respect to states of India. With bar plot deprecating numbers of cases per states.

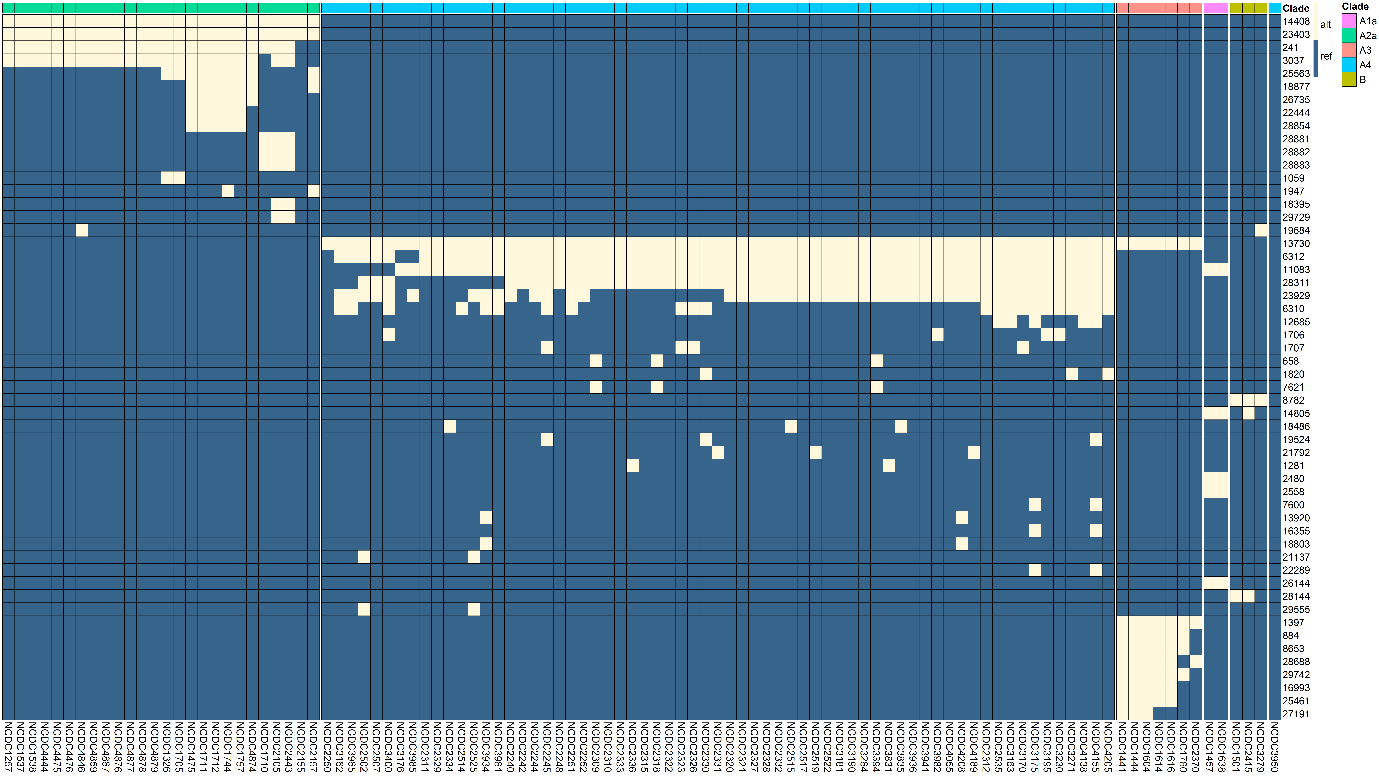

**Figure-S2**: a heat map representation of SARS-CoV-2 variants per sample and their respective segregation in respective clusters. The light pink colour indicates presence of mutation and background blue indicates wild type allele.

Top bar at each cluster indicates respective clade information.
